# ploidyNGS: Visually exploring ploidy with Next Generation Sequencing data

**DOI:** 10.1101/086488

**Authors:** Renato Augusto Corrêa dos Santos, Gustavo Henrique Goldman, Diego Mauricio Riaño-Pachón

## Abstract

**Summary:** `ploidyNGS` is a model-free, open source tool to visualize and explore ploidy levels in a newly sequenced genome, exploiting short read data. We tested `ploidyNGS` using both simulated and real NGS data of the model yeast *Saccharomyces cerevisiae*. `ploidyNGS` allows the identification of the ploidy level of a newly sequenced genome in a visual way.

**Availability and implementation:** `ploidyNGS` is available under the GNU General Public License (GPL) at https://github.com/diriano/ploidyNGS. `ploidyNGS` is implemented in Python and R.

## 1 Introduction

Knowledge about the ploidy level of an organism or cell type is an important piece of information in evolutionary, population and genomic studies. Ploidy level information can usually be obtained with laborious flow cytometry experiments [Kron *et al*., 2007]. However, as the costs of Next Generation Sequencing (NGS) decrease, and data from Whole Genome Sequencing (WGS) projects become increasingly available, tools that exploit short-read sequencing data to infer ploidy levels are being increasingly sought after.

Some methods have recently been developed for ploidy estimation using NGS data. AbsCN-seq [Bao *et al*., 2014] and CLImAT [Yu *et al*., 2014] were developed with the purpose of analyzing tumor samples, and both include ploidy detection. However, absCN-seq requires additional information besides mapped sequencing reads as well as Whole Exome Sequencing data, while CLImAT is based on MATLAB, which is not freely available. ConPADE [Margarido and Heckerman, 2015] was developed for the estimation of ploidy in highly polyploid plant genomes; however, its underlying model is sensitive to the quality of the mapping step, which may cause an upward bias in ploidy estimates (personal communication from the author).

One can get a good sense of the ploidy level by counting the number of reads supporting different alleles at each position along the genome sequence. For instance, in a haploid organism, at every position along the genome, all reads will support a single allele, excluding sequencing errors. In a diploid organism, depending on the heterozigosity level, in addition to monomorphic positions one would find positions where half of the reads support one allele and the other half support an alternate allele. In the case of a triploid, in multiallelic sites a third of the reads will support each allele, in bi-allelic sites, 2/3 of the reads would support one allele and the remainder third would support the other one, and so on for further ploidy levels [Ludlow *et al*., 2016, Yoshida *et al*., 2013]. Hence, looking at the frequency of the different observed alleles offers a picture of the ploidy level. We have automated this process in `ploidyNGS`, to study the frequency profiles for each putative allele in a histogram. For instance, in a diploid, in heteromorphic positions, the most frequent allele has a frequency close to 50%, as well as the second most frequent allele, so we expect a peak close to 50% with a height that depends on the heterozigosity level. We tested the tool using several simulated *Saccharomyces cerevisiae* autopolyploid genomes, with different ploidy and heterozigosity levels, generating reads from these simulated genomes using the error profile of the Illumina chemistry [Huang *et al*., 2011]. In addition, we tested our software on real WGS data of known haploid and diploid *S*. *cerevisiae* strains available in the Saccharomyces Genome Database (SGD, http://www.yeastgenome.org/).

## 2 Methods

### 2.1 Description of the algorithm

The script `explorePloidyNGS.py`, does all the computation, and it requires as input a BAM file, with the mapping of genomic reads to the genome sequence. The BAM file can be easily generated using tools such as bowtie2 [Langmead and Salzberg, 2012]. `ploidyNGS` processes this file in the following way:

- It does a first pass on the file, storing, in a data structure, for each position, the number of reads supporting different nucleotides, i.e., putative allele abundance
- Then it traverses the data structure, ignoring positions where a single nucleotide was observed and where the most frequent nucleotide had a frequency larger than the parameter `max_allele_freq`, which is set by the user (default 0.95). For the remaining positions, it computes the putative allele percentage as the fraction of reads supporting a given nucleotide over the sequencing depth at the given position.
- Putative allele percentages at each position are ordered from lowest to highest, and labeled as fourth, third, second and first.
- Finally, a histogram is generated using the ggplot2 package in R, for all the labels generated in the previous step. This histogram can be compared to the results of simulated datasets available in the github repository (https://github.com/diriano/ploidyNG/tree/master/simulation/results) to help on deciding which is the possible ploidy level of the organism under study.

## 3 Results

### 3.1 Simulated data

We have generated simulated data, representing different ploidy (2 to 7) and heterozygosity (10^-1^ to 10^-4^) levels, as well as different sequencing depths (10x, 25x, 50x and 100x), as a reference for comparisons. All the simulated datasets were derived from the *S*. *cerevisiae* S288C genome sequence. Autopolyploid genomes were generated with the script `simulatePloidyData.py` (a detailed description of this script is available at the `ploidyNGS` GitHub repository).

Sequencing depth was simulated by generating short paired-end reads mimicking the error profile observed for Illumina [Huang *et al*., 2011]. For each combination of ploidy level, heterozigosity and sequencing depth, mapping files were generated using bowtie2 with default parameters [Langmead and Salzberg, 2012]. Results for all the simulated datasets are available in the GitHub repository, and can be used as aid to assess ploidy levels for new organisms. The visual identification of the ploidy level largely depends on the proportion of heteromorphic positions, and the sequencing depth.

### 3.2 Real data

We applied `ploidyNGS` to real yeast genome datasets obtained from SGD, for which the ploidy were previously known (Figure 1). In the case of the haploid strain (Figure 1A), the most frequent allele is close to 95% (higher frequencies were excluded with the parameter `max_allele_freq`, the second most frequent allele has an abundance of less than 5% and most likely represent sequencing errors. In Figure 1B, the most frequent allele has a peak for monomorphic positions, close to 95%, and a peak close to 50% for heterozygous positions, and the second most frequent allele has a peak close to 50%, which represent heterozygous positions, and another one close to 5% which represent sequencing errors, hence compatible with a diplod genome. Reads for both strains were obtained from SRA (SRR445715 and SRR1569660).

**Figure 1:**
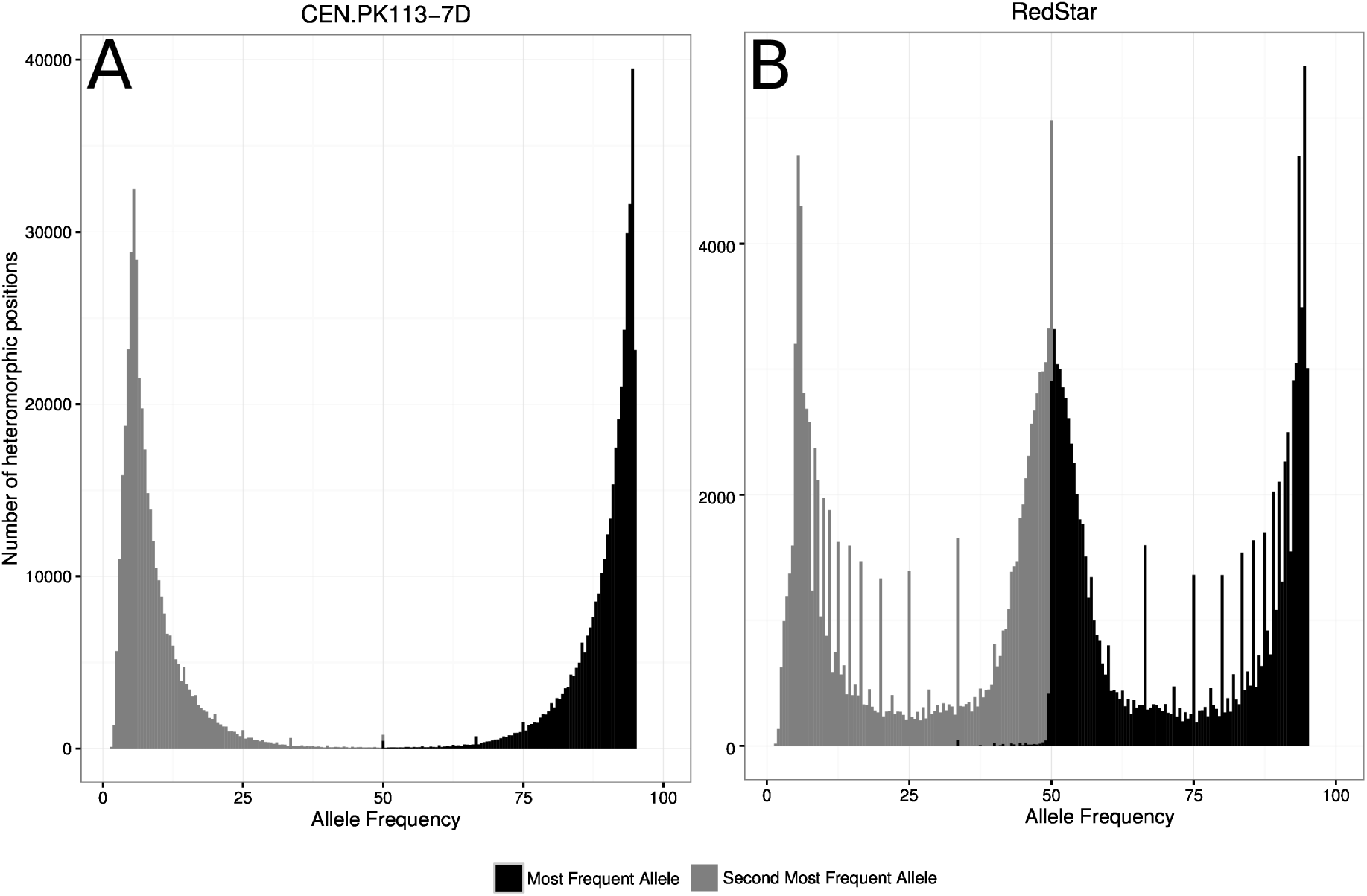
Exemplary results from `ploidyNGS`, here just showing the two most frequent putative alleles A) Haploid yeast strain CEN.PK113-7D B) Diploid yeast strain RedStart.

## 4 Conclusion

`ploidyNGS` is a useful tool that allows to visually assess the ploidy of organisms for which there is NGS short-read data available. It has been tested on eukaryotes with small genome sizes such as yeasts, filamentous fungi and oomycetes.

## References

Bao, L., Pu, M., and Messer, K. (2014). AbsCN-seq: a statistical method to estimate tumor purity, ploidy and absolute copy numbers from next-generation sequencing data. Bioinformatics, 30(8), 1056–1063.

Huang, W., Li, L., Myers, J. R., and Marth, G. T. (2011). ART: a next-generation sequencing read simulator. Bioinformatics, 28(4), 593–594.

Kron, P., Suda, J., and Husband, B. C. (2007). Applications of flow cytometry to evolutionary and population biology. Annu. Rev. Ecol. Evol. Syst., 38(1), 847–876.

Langmead, B. and Salzberg, S. L. (2012). Fast gapped-read alignment with bowtie 2. Nature Methods, 9(4), 357–359.

Ludlow, C. L., Cromie, G. A., Garmendia-Torres, C., Sirr, A., Hays, M., Field, C., Jeffery, E. W., Fay, J. C., and Dudley, A. M. (2016). Independent origins of yeast associated with coffee and cacao fermentation. Curr. Biol., 26(7), 965–971.

Margarido, G. R. A. and Heckerman, D. (2015). ConPADE: Genome assembly ploidy estimation from next-generation sequencing data. PLoS Comput. Biol., 11(4), e1004229.

Yoshida, K., Schuenemann, V. J., Cano, L. M., Pais, M., Mishra, B., Sharma, R., Lanz, C., Martin, F. N., Kamoun, S., Krause, J., Thines, M., Weigel, D., and Burbano, H. A. (2013). The rise and fall of the *Phytophthora infestans* lineage that triggered the irish potato famine. eLife, 2.

Yu, Z., Liu, Y., Shen, Y., Wang, M., and Li, A. (2014). CLImAT: accurate detection of copy number alteration and loss of heterozygosity in impure and aneuploid tumor samples using whole-genome sequencing data. Bioinformatics, 30(18), 2576–2583.

